# Engineering Endogenous T Cell Receptors to Recognize Cancer Neoantigens Using a Hybrid Physics-AI Approach

**DOI:** 10.64898/2026.05.15.725176

**Authors:** Jeffrey K. Weber, Gyanu Parajuli, Stephen Wang, Vadim Ratner, Xiaoxiao Ma, Yoel Shoshan, Leili Zhang, Joseph A. Morrone, Moshiko Raboh, Efrat Hexter, Prerana Parthasarathy, Paul Pavicic, Christina Gaughan, Vladimir Makarov, Lawrence Chu, Suheyla Hasgur, Ivan Juric, C. Marcela Diaz-Montero, Raghvendra Srivastava, Jeffrey A. Knauf, Khaled A. Hassan, Wendy D. Cornell, Tyler J. Alban, Timothy A. Chan

## Abstract

T cell receptors (TCRs) are critical for immune surveillance and successful adaptive immune response against foreign antigens. TCRs drive this key arm of the immune system through recognition of peptide epitopes presented on MHC complexes. However, they are limited due to their stochastic nature and generation via genetic recombination. In silico design of functional TCRs that target defined peptide epitopes would be of considerable utility but has up until now been unsuccessful. Here, we develop an artificial intelligence (AI)-powered approach using a hybrid physics-based simulation and generative AI that successfully engineers TCRs against defined epitopes presented by MHC-I. We use this approach to design TCRs against two cancer antigens, a HERC1 neoantigen and an immunogenic neoepitope in mutant EGFR. We engineer multiple TCRs against the HERC1 neoantigen which activate T cells in response to exposure to peptide-MHC I and kill cancer cells more effectively than a patient-derived TCR. In addition, we used generative AI to design functional TCRs that target the EGFR T790M neoantigen, engineering greater specificity against the mutant sequence. We present an AI-based approach to TCR design with broad utility for efforts to engineer TCRs and for the development of new cell therapies.

**One sentence summary:** Artificial intelligence-based approach enables the directed engineering of functional TCRs with enhanced features that target cancer neoantigens.

## Introduction

Major histocompatibility complex (MHC) class 1 molecules present short peptides on the surfaces of cells, enabling cytotoxic T cells to continuously survey the MHC peptidome for foreign proteins. This process is critical for the ability of the immune system to act against cancerous cells or pathogen. In humans, these molecules are encoded by highly polymorphic human leukocyte antigen (HLA) genes, which dictate peptide presentation and shape immune recognition. Cancer neoantigens, which often differ from self-peptides by a single point mutation, represent attractive targets for immunotherapy (*1-4*). However, achieving selective recognition of mutant peptides while avoiding wild-type counterparts demands exceptional specificity. In addition to specificity, endogenous TCRs must engage MHC with affinities tuned for function, balancing sufficient activation with reduced risk of cross-reactivity (*5, 6*). These criteria are met by the immune system through stochastic generation of TCRs and a series of positive and negative selections, ultimately resulting in functional TCRs. However, stochasticity, immunologic health, age, and T cell repertoire holes can limit the spectrum of TCR diversity.

In contrast, biologics such as antibodies undergo affinity maturation via somatic hypermutation (*7*). After recombination, the immune system refines their affinity to improve affinity. This mechanism has aided development of biologics such as antibodies, nanobodies, and engineered scaffolds for peptide MHC (pMHC) complex targeting (*8*). While effective in some contexts, these engineered non-native molecules introduce therapeutic liabilities, such as unpredictable off-target binding, immunological exhaustion, and degradation challenges. Lengthy and unreliable *ad hoc* screening methods for TCR discovery have been the primary option for TCR development. Engineering cytotoxic T cell receptors (TCRs) in their natural αβ format offers a physiologically aligned approach, but the exact rules defining TCR recognition of pMHC have not been fully resolved.

To approach this problem, recent advancements in generative AI, LLMs, and diffusion-based artificial intelligence (AI) architectures have been used to design protein-based therapeutics (*2, 9-20*). Multiple examples have emerged including the design of helix bundles that can be grafted onto CAR-T cell architectures and the creation of functional antibodies and nanobodies. Unlike antibodies, TCRs do not undergo affinity maturation and engineered TCR expressing T cells mimic the natural pathway for class I antigen-specific cytotoxicity. TCRs from natural T cells can have a range of affinities, varying abilities to activate T cells, and engagement of downstream effector functions. Therefore, the ability to engineer TCRs would be of substantial benefit for efforts to develop TCR based cell therapies.

Here, we report results of an *in-silico* approach to engineering functional cytotoxic TCRs in their natural format. Our pipeline combines physics-based simulations, which provide atomistic insight into interactions at the class I MHC/peptide–TCR interface, with multiple AI models built on graph-based and large language model (LLM) architectures. As our first use case, we focused on one of the first human neoantigens described, a HECR1 neoantigen (*21*). Targeting this HERC1 mutation (P3278S, ASNA**S**SAAK), our approach yielded a field of candidates. We took forward 8 candidates to test functionally. Two functional TCRs out of eight designs (25%) were capable of both pMHC targeting and tumor cell killing. Consistent with their design criteria, these engineered candidates demonstrated stronger activation and cytotoxicity compared to a TCR identified from patients. Building on this success, we next focused on the EGFR neoantigen T790M (**M**QLMPFGCLL)(*22*). Using our AI strategy, we developed functional TCRs that recognize the EGFR T790M neoepitope, with one engineered candidate with increased specificity to mutant peptide.

## Results

We developed a combinatorial AI workflow for TCR design. This system integrates free-energy perturbation (FEP) simulations (*23-28*) with AI-driven sequence generation and ranking to optimize TCRs for relative affinity and specificity. FEP provides atomistic insight into binding energetics at the pMHC–TCR interface, enabling accurate prediction of favorable CDR3 regions, while AI models, including graph-based predictors and large language models (LLMs), rapidly prioritize candidates and explore sequence space beyond what might be experimentally tractable (**Fig. 1A**). This synchronization of physics-based free energy simulations (FEP) and LLMs provides unique advantages to previous TCR design approaches, as knowledge of the massive corpus of known TCR sequences can be blended with accurate models of atomistic interactions between highly variable TCR CDRs and presented class I peptides. Experience from this hybrid approach was further used to inform the development of pure LLM methods for generating TCR candidates, as described below and in the Supplementary Methods.

**Fig. 1.**
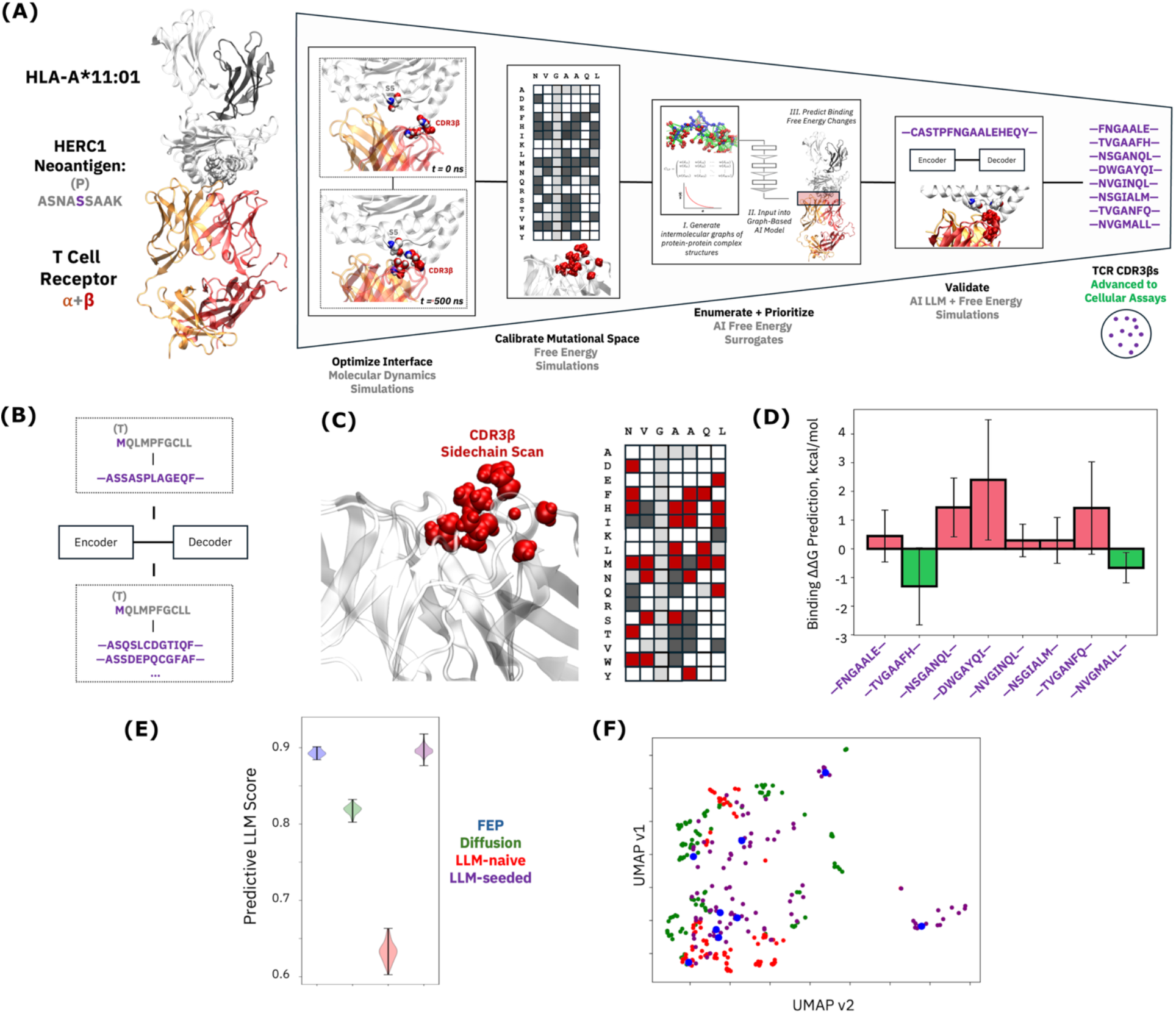
Design of TCRs with an artificial intelligence (AI)-powered approach using hybrid physics-based simulation and generative AI. (A) Ternary protein complex (TCR-neoantigen-MHC) for cancer neoantigen recognition by cytotoxic T cells that feeds into hybrid simulation and AI workflow for engineering TCR candidates. This workflow is based on molecular dynamics and free energy simulations as well as sequence (LLM)- and structure-based AI models. The HERC1 protein neoantigen (ASNASSAAK) is shown. (B) Schematic of TCR candidate generation and validation pipeline with LLM-based AI model. (C) Results of point mutational scan of patient-derived TCR for HERC1 neoantigen binding. Red squares indicate mutations for which improved binding (negative ΔΔG) is predicted and that were advanced to candidate enumeration with FEP surrogate models. Dark grey squares represent predicted binding free energy differences in [0,1] kcal/mol, and white squares > 1 kcal/mol; values for light grey squares were not computed. (D) Predictions of binding free energy changes of final eight TCR candidates advanced to cellular assays, with 95% confidence intervals. (E) Violin plot of predictive LLM scores for TCR candidates drawn from four generative classes. Probabilities are estimated with softmax normalization. (F) UMAP projection of circular fingerprints of TCR CDR3β sequences drawn from four generative classes.

In combination with FEP, we applied both graph- and LLM-based models specifically to TCR candidate prioritization and an LLM-based approach to candidate generation. Graph-based models were founded on graph convolutional architectures classifying MHC-peptide complexes (*29, 30*). LLMs were based on a novel, multi-modal foundation model (MAMMAL) that has been specifically optimized for predicting interactions between biological macromolecules (*31*). This model, pretrained on approximately 2 billion diverse examples including TCRs and protein-protein interactions, employs a structured prompt syntax that allows direct encoding of TCR-epitope complexes. MAMMAL demonstrates superior performance on TCR-epitope binding tasks, improving upon previous sequence-based methods without relying on structural information (**Fig. 1A; Supplementary Methods**) (*32-35*).

### In silico engineering of TCRs targeting HERC1 neoantigen

We aimed to design TCRs that had superior features compared to ones that we identified using cell-based screens from patient materials. We identified a patient-derived TCR that targets MHC-bound HERC1 neoantigen (MT: ASNA**S**SAAK; WT: ASNA**P**SAAK; HLA-A*11:01)(**Fig. 1A**), demonstrating that T cells can target this neoantigen when presented by MHC I. To engineer the CDR3β to increase activation, we conducted exhaustive free-energy perturbation (FEP) simulations of every possible point mutant within the central, seven-residue portion of the reference TCR CDR3β (-NVGAAQL-). From this analysis, we identified substitutions likely to enhance binding (**Fig. 1A**). Substitutions predicted to enhance binding were combined into double, triple, and quadruple) mutants, which were then constructed through all possible linear combinations of non-destructive point mutations. These combinations were subsequently ranked using our graph-based AI model trained on single-mutant FEP data. Eight candidates were advanced to full FEP simulations and, in parallel, scored by our LLM fine-tuned for pMHC–TCR binding prediction (**Fig. 1A**). Sequences were also generated and ranked using pure iterative AI approaches (**Fig. 1B**).

The mutational perturbations of HERC1 TCR CDR3β positions revealed 28 substitutions that were predicted to improve binding affinity to HERC1 neoantigen ASNA**S**SAAK in isolation, with diverse interchanges of polar, hydrophobic, and charged residues at each position (**Fig. 1C**). Multiple mutant constructs were ranked and clustered using a graph-based FEP surrogate AI model trained on free energy simulation data. **Fig. 1D** shows the ΔΔG from FEP simulations of the multiple mutant systems. Two candidates (--TVGAAFH-- and –NVGMALL--) were predicted to bind more strongly than the patient TCR, with all eight sequences showing promise at 95% confidence.

Our LLM fine-tuned on TCR sequence data confirmed the previous prioritization; all eight sequences scoring at an approximately 90% binding probability, compared to an average of ∼65% for candidates naively generated by the LLM (**Fig. 1E**). Sequences generated by an LLM seeded with the hybrid workflow candidates recovered, or even exceeded, binding probabilities of the original TCRs. Interestingly, diffusion-generated sequences (*20, 36, 37*) were predicted to bind at levels above the naïve LLM candidates but remained below the hybrid and seeded-LLM designs. UMAP projections (*38*) revealed reasonably distinct sequence clusters from each design method (**Fig. 1F**), with seeded LLM sequences sampling neighborhoods falling closest to the hybrid workflow candidates. **Figures**

### Functional validation of engineered TCRs targeting the HERC1 neoantigen

Eight TCR candidates engineered by the pipeline to target the HERC1 P3278S neoantigen (ASNASSAAK, HLA-A*11:01) were optimized for expression and examined using Jurkat reporter assays and primary CD8^+^T cell cytotoxicity assays (**Fig. 2A, B**). Out of the 8 TCR candidates that we moved forward, 8/8 (100%) made mature TCRs that localized to the T cell surface (**Supplemental Fig. 1A**). Two out of the eight engineered TCRs (25%), EC_2 and EC_8, showed TCR signaling activation using the NFAT reporter assay with mutant peptide presenting APCs as target cells (**Fig. 2C**). Consistent with the NFAT reporter activation, we also noted that EC_2 and EC_8 had increased induction of activation markers including CD69 and PD-1 (**Fig. 2D, Supplemental Fig. 1B**). Furthermore, TCRs EC_2 and EC_8 retained mutant peptide specificity with minimal cross reactivity to wild-type HERC1 peptide (**Fig. 2E**). Toxicity related cytokine TNFα also demonstrated mutant peptide specificity for EC_2 and EC_8 (**Fig. 2F**).

**Fig 2.**
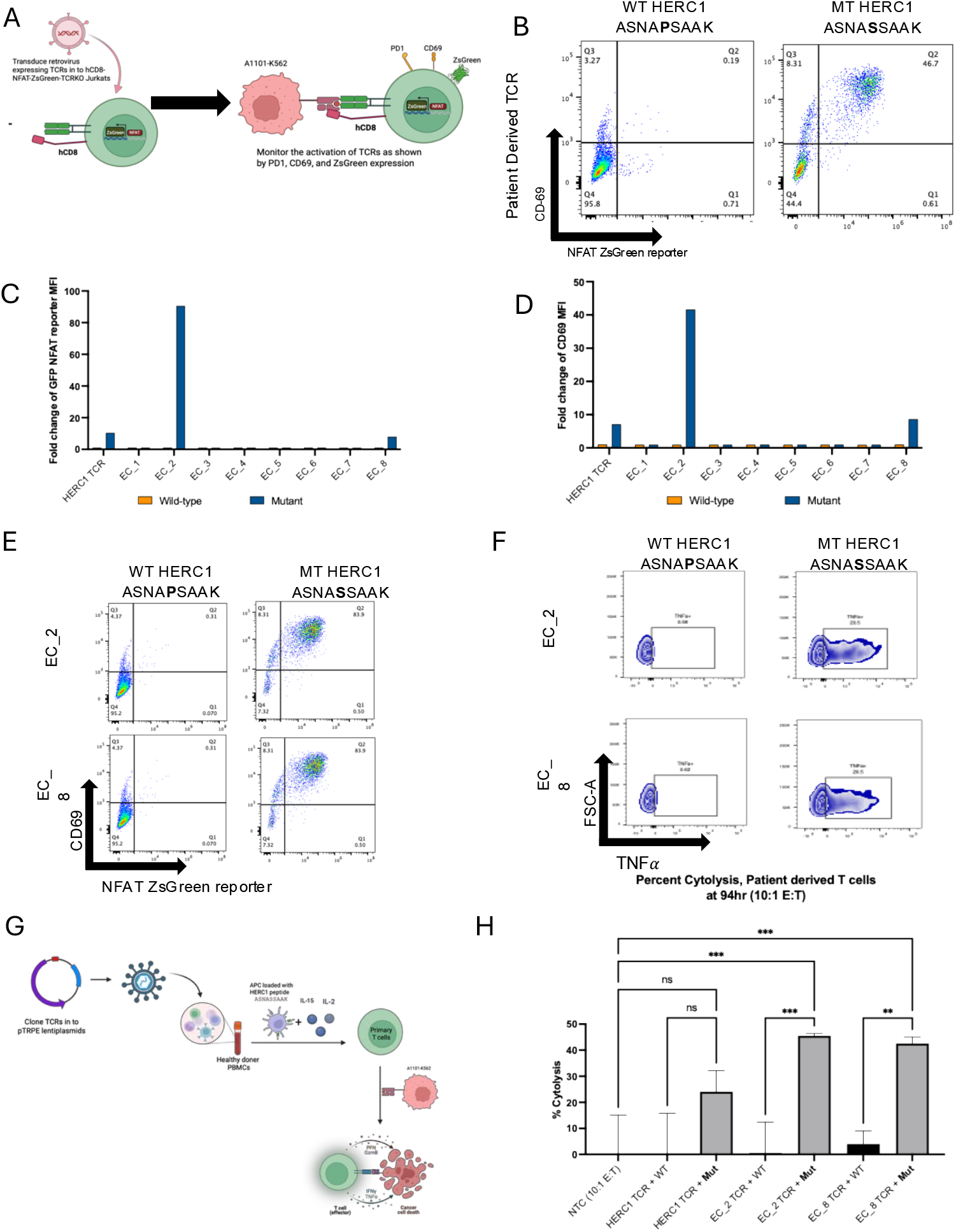
Mutant specific activation and cell killing by designed TCRs targeting HERC1. (A) Illustration showing the workflow of TCR activation assay. (B-F), hCD8-NFAT-Jurkats were transduced with HERC1 TCRs and AI-designed TCRs. Activation of HERC1 TCRS by K562 cells expressing HLA-A*11:01 pulsed with 10 μM of WT HERC1 (ASNAPSAAK) and Mutant (MT) HERC1 (ASNASSAAK) as shown by NFAT ZsGreen reporter+CD69+ expression (B,E), NFAT ZsGreen reporter expression (C), and CD69 expression (D). MFI of protein was normalized to DMSO control to calculate fold change. (F) Expression of TNF α in AI-Designed TCRs when incubated with HLA-A*11:01-K562s pulsed with 10 μM of WT and MT HERC1 peptides. (G) Illustration showing the mutant specific cytotoxicity by HERC1 TCRS. (H) Primary T cells were transduced to express HERC1 TCRs and AI-derived TCRS, and co-cultured with HLA-A*11:01-K562s. T cell–mediated cytolysis was monitored using the xCELLigence Real-Time Cell Analyzer. Cell indices plotted in the bar graphs represent the mean of triplicate samples, error bars represent standard deviation and *p-*values calculated from one-way ANOVA with Tukey’s multiple comparison test (*=p<0.05, **=p<0.01, ***=p<0.001). The result shown is representative of 2 independent experiments. NTC, non-transduced T cells.

With clear reporter T cell activation by two candidates, we next sought to evaluate the engineered TCR candidates’ ability to kill target cells using primary T cells. **Fig. 2G** shows our experimental design. T cells expressing the EC_2 or EC_8 engineered TCRs killed HERC1 mutant presenting APCs efficiently compared to controls (**Fig. 2H**). Furthermore, whereas the original patient derived HERC1 TCR was able to activate reporter, it did not enable the effective killing of neoantigen-presenting antigen presenting cells compared to the non-target, empty vector control (**Fig. 2H**). Time courses of T cell killing showed that the patient derived TCR does demonstrate some cytotoxicity, but the killing activity was relatively weak (**Supplemental Fig. 1C**). In contrast, AI-designed and optimized TCRs EC_2 and EC_8 demonstrated superior cytotoxicity compared to the patient derived TCR (**Supplemental Fig. 1 D, E**). Furthermore, EC_2 and EC_8 also showed significant specificity, killing mutant HERC1 peptide presenting target cells but not the wild-type peptide presenting APCs (**Fig. 2H**).

### FEP naïve engineering approach for EGFR T790M targeted TCRs

To further examine the utility of our generative AI model, we applied our LLM alone-based TCR pipeline (without molecular dynamics) to the EGFR T790M neoantigen (wild-type: **T**QLMPFGCLL; mutant: **M**QLMPFGCLL HLA-A*02:05). This neoantigen is a particularly challenging use case as the mutation is located at position 1 of the neopeptide. Amino acids in this position lie in the minor groove of the MHC molecule and do not face outward along the MHC major groove. Candidates were generated using both naïve and iterative LLM approaches (see SI). This neoantigen represents a significant challenge since the mutation occurs at the peptide terminus rather than the central TCR contact region, making discrimination between mutant and wild-type peptides particularly difficult. Attempts at identifying TCRs using donor derived T cells in an extensive ad hoc screening attempt yielded 3 clones that recognized both MT and WT peptides, underscoring the complexity of achieving true mutant specificity. In this case, we aimed to use in silico design to engineer TCRs capable of recognizing greater the mutant peptide with greater selectivity. Using this approach, 20 candidates were identified to have predicted ΔΔG more favorable to binding MHC-presented mutant peptide vs wild-type peptide.

We examined the function of the AI engineered TCRs. Each of the 20 TCR candidates were cloned into Jurkat cells with the NFAT reporter as above. Out of the 20 EGFR TCR candidates that we moved forward, 18 (90%) made mature TCRs that localized to the T cell surface (**Supplemental Fig. 2A-C**). Two of these candidates, TCR EC_7-7, and EC_750-5, were able to activate NFAT in response to mutant EGFR neoantigen peptide (10%)(**Fig. 3A**). Furthermore, these candidates also induced CD69 expression (**Fig. 3B**), but only EC_750-5 led to robust induction of PD1 expression (**Supplemental Fig. 2D**). To evaluate how specific the newly engineered clones were to mutant versus wild-type neopeptide compared to the patient derived TCR, we plotted the fold change in activation of the difference between wild-type peptide stimulated activation and mutant peptide stimulated activation. The donor derived TCR 7 had significantly stronger activation from wild-type peptide compared to mutant, which was not seen in the engineered EC_7-7 TCR (**Fig. 3C**). Similarly, donor derived TCR 750 also showed significantly more wild-type targeting than mutant. But, strikingly, the engineered TCR EC_750-5 had significantly increased mutant specificity (**Fig. 3D**).

**Figure 3.**
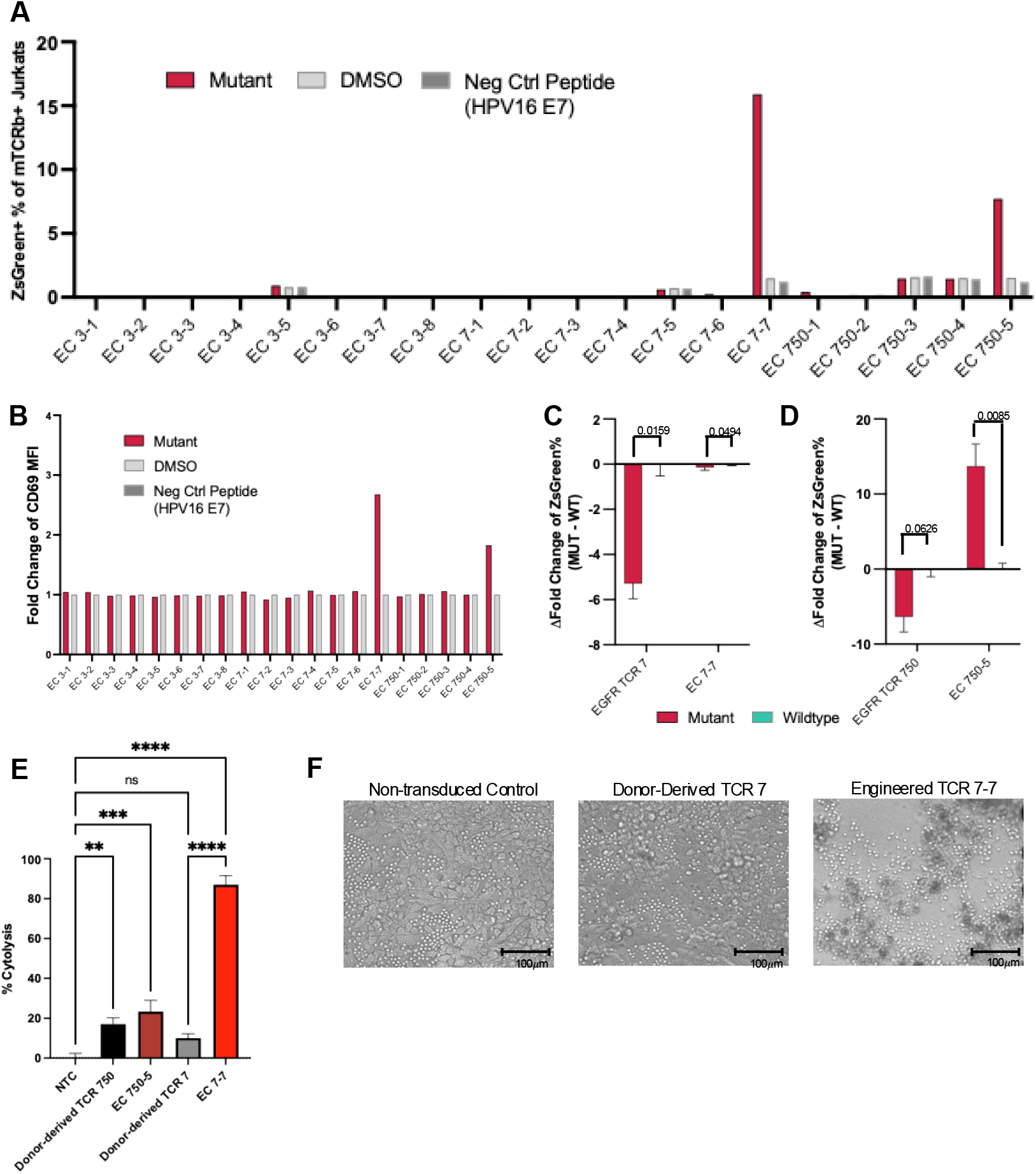
Identification and characterization of engineered TCRs targeting the EGFR T790M neoepitope. (A) Jurkat reporter activation assay was performed for 20 engineered TCRs. Displayed is the percent of engineered TCR-expressing Jurkats that are ZsGreen-positive in response to the T790M neoepitope, DMSO vehicle control, and a negative control peptide from HPV16 E7. (B) Quantification of CD69 surface expression MFI in response to T790M neopeptide was normalized to DMSO controls to calculate fold change. (C) Change in activation between mutant peptide stimulation minus wild-type peptide stimulation for donor-derived TCR 7 and EC_7-7. (D) Change in activation between mutant peptide stimulation minus wild-type peptide stimulation for donor derived TCR 750 and EC_750-5. For C&D, the more positive the value, the more mutant-specificity is demonstrated by the TCR. Fold change of ZsGreen reporter positivity relative to DMSO was calculated and wildtype subtracted from mutant. Error bars represent standard deviation and *p-*values calculated from unpaired t-test. (E) Cytotoxicity assay using H1975 A*02:05 tumor cell line that natively expresses mutant EGFR T790M and expresses HLA-A*02:05. Error bars represent standard deviation and *p-*values calculated from one-way ANOVA with Tukey’s multiple comparison test (*=p<0.05, **=p<0.01, ***=p<0.001). (F) Representative images at of non-transduced T cell control, TCR 7 and EC 7_7 at 48hrs after T cell addition. Images acquired at 10x magnification with phase-contrast microscope.

To evaluate the ability of the engineered TCRs to promote T cell cytotoxicity, we examined their ability to kill EGFR T790M mutant gene expressing tumor cells. In this assay, the patient derived tumor cell line H1975, which harbors the EGFR T90M mutation, was cultured with primary T cells expressing the engineered TCRs. While primary T cells expressing TCR 750 demonstrated cytotoxic activity, donor derived TCR 7 did not (**Fig. 3E**). Strikingly, TCR EC_7-7 had a 9-fold increase in killing compared to donor derived TCR 7 (**Fig. 3E**). **Figure 3F** shows example images of tumor cell killing by T cells expressing the engineered TCRs.

### Structural modeling reveals mechanisms associated with engineered TCR function

With successful generation of TCRs that activated NFAT reporter activity and killed targets cells by targeting neopeptide-MHC, we next sought to understand the features associated with the successful AI-driven designs. The physics-based simulations conducted to prioritize initial HERC1 candidates provide a window into binding mechanisms for the two HERC1 TCR hits and contextualized potential interaction differences relative to the patient derived reference TCR. Overall, the engineered CDR3β sequences appear to benefit from both enthalpic and entropic advantages to binding. While the patient-derived TCR is predicted to favor interactions between the mutant position in the neoantigen (S5) and the C-terminus of the CDR3β (**Fig. 4A**), engineered TCR sequences are predicted to engage via a register shift that integrates the full CDR3β loop, allowing for additional interactions at the N-terminus that could contribute to TCR-peptide binding. No stable interactions are observed between the wildtype peptide (which is drawn into the MHC peptide-binding groove in proline-favored presentations) and the patient-derived CDR3β, perhaps explaining the resilience of both reference and engineered TCRs in avoiding wildtype activation. We also noted that in HERC1 engineered TCRs, changes introduced appear to reduce the conformational flexibility of the TCR CDR3βs in their free states (**Fig. 4B**), a factor that would generally reduce the entropic cost to binding the cancer neoantigen in a process that fixes the loop in place. Central and C-terminal regions of the loop, at the edge and outside of the loop helix that has helical propensity, seem to be more flexible in the reference TCR. The combination of increased interactive regions and lesser loss of flexibility seem to enhance binding affinity in the engineered TCRs.

**Fig. 4.**
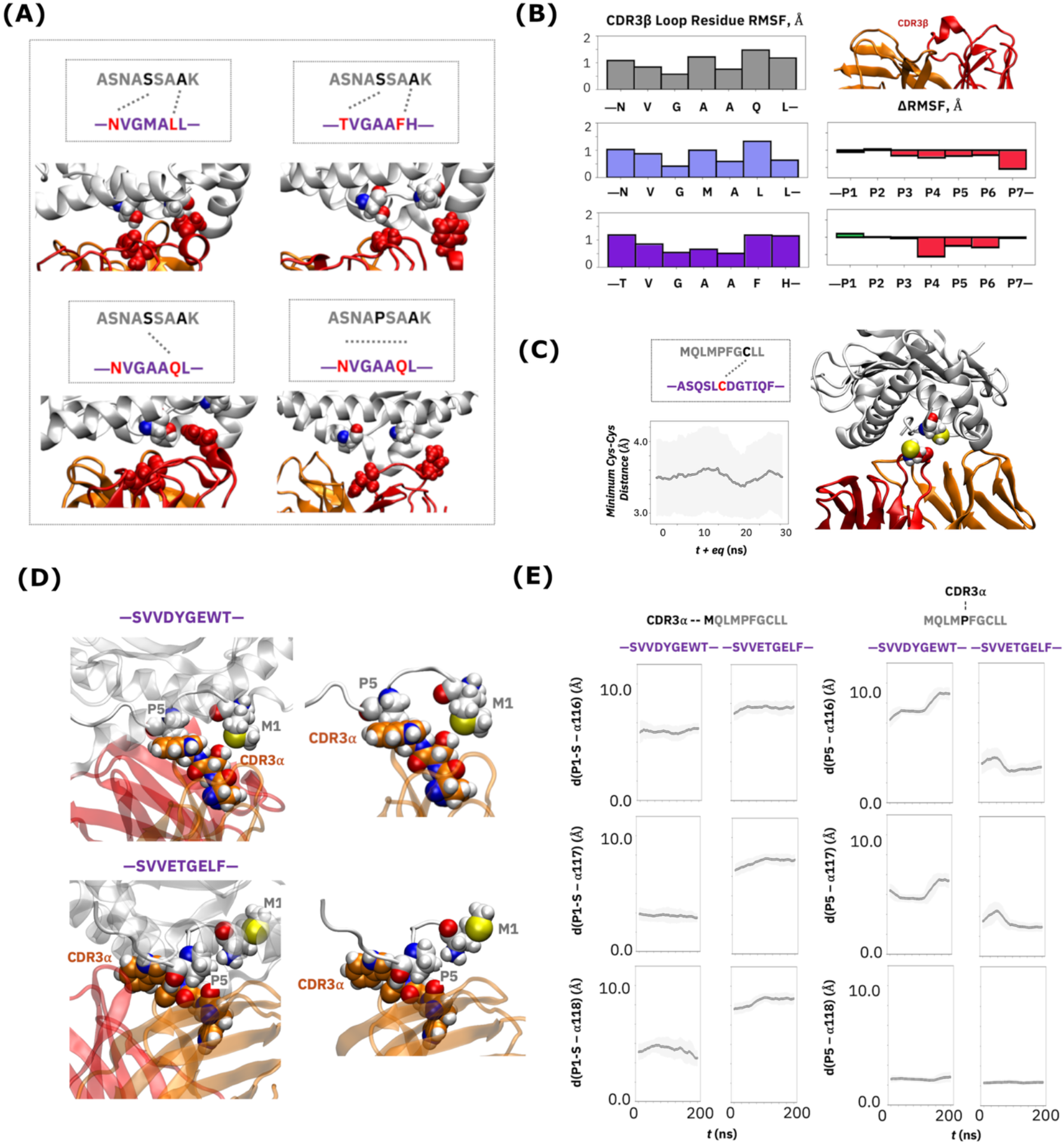
Structural and physicochemical analysis reveals mechanisms of experimentally validated engineered TCRs. (A) Comparison of predicted interactions between the HERC1 cancer neoantigen and *in-silico* and patient-derived TCRs that suggest mechanisms behind binding enhancements. Simulations show minimal interactions between the patient-derived TCR CDR3β and the wild-type peptide. (B) TCR CDR3β loop root mean square fluctuations (RMSFs) for patient-derived and in-silico activators against the HERC1 neoantigen, along with RMSF differences between candidates and the patient-derived reference. (C) TCR-peptide interactions between EC_7-7 against EGFR T790M neoantigen, highlighting the proximity of cysteine residues. (D) Comparison of putative binding modes between 750-5 TCR candidate and patient-derived TCR sequence showing differential neoantigen-TCR interactions at the mutational locus (P1: T2M). (E) CDR3-peptide minimum distance plots quantifying the conformational and interactive changes illustrated by structural renderings.

Developing TCRs that specifically target EGFR T790M neopeptide-MHC was, as expected, more challenging. Here, we interrogated the AI-driven mechanisms of successful designs. The C7-7 TCR candidate had strong killing properties but lacked specificity against the mutant, which was similar to the patient derived TCR. One intriguing possibility may be related to a disulfide bonding mechanism for activation (**Fig. 4C**). Although cysteines in TCR CDR3s are not as common as TCR CDR3 without cysteines, the presence of cysteines in TCRs can promote strong T cell activation(*39*) and inclusion of cysteines have been used to engineer affinity-enhanced TCRs(*40, 41*). MD simulations of the ternary pMHC-TCR candidate complex show a strong propensity for the cysteine residues on either side of the synapse to colocalize, with minimum distances rarely exceeding 4Å (**Fig. 4C**). Based on this, we revisited the prediction frequency of cysteines and found that EGFR predicted CDR3B sequences were more likely to have a cysteine than in the HERC1 predictions (**Supplemental Fig. 3A**). To understand how much this conceivable mechanism contributes to activation, we generated mutant peptides C8S, C8A, C8G and an experimentally verified 9mer from another publicly available dataset that is HLA-A*02:05 restricted and has a cysteine (*42*). We first experimentally confirmed that all peptides bound to HLA-A*02:05 (**Supplemental Fig. 3B, C**) and then examined TCR activation profiles. Assays demonstrated that EC7_7 was activated by mutant and wild-type EGFR peptides, but in both cases lost activation with the absence of cysteine (**Supplemental Fig. 3D**). Importantly, the unrelated 9mer sequence with cysteine did not cause non-specific association (via disulfide bonding of EC_7-7 to the T790M peptide) (**Supplemental Fig. 3D**).

In contrast, TCR EC_750-5 demonstrated more moderate activation and killing but showed a significant increase in specificity compared to the patient derived TCR. Structural modeling showed that the patient-derived TCR is predicted to favor interactions between the mutant position in the neoantigen (S5) and the C-terminus of the CDR3β, while EC7_750-5 showed preference towards the mutated M1 position (**Fig. 4D**). MD simulations further suggest a putative mechanism for neoantigen binding specificity observed with the EC_750-5 TCR candidate. The introduction of bulky tyrosine and tryptophan residues to the CDR3β sequences appears to shift the register of interactions between the peptide and the TCR CDR3α. While the reference sequence favors contacts between the center of the neoantigen, particularly with the P5 proline, CDR3α contacts of the TCR 750-5 candidate are dominated by the C-terminal methionine of the neoantigen that represents the cancer mutation in this system (**Fig. 4D**). That induced dependence of neoantigen interactions on the cancer mutational locus likely leads to the neoantigen specificity observed in experimental screens. Minimum distance plots between the neoantigen and CDR3α quantify the interaction differences, with M1 sulfur-CDR contacts emerging and P5-CDR candidates disappearing in the C750-5 candidate (**Fig. 4E**).

## Discussion

Antibodies can be engineered to target virtually any extracellular epitope, an attribute that has rendered them highly versatile. By contrast, classic T cell receptors (TCRs), which form the other arm of the adaptive immune system, specialize in recognizing antigens presented by MHC molecules and serve as a critical mechanism for continuous immune surveillance. Despite their immense therapeutic potential, TCRs have not been fully leveraged in clinical applications and lack of affinity maturation limits the scope of potential biological solutions. The need for TCR-based therapeutics is substantial, yet their development has been hindered by the absence of an efficient means for precision design.

Our results demonstrate that an AI-guided engineering approach, alone or hybridized with physics-based simulations, can generate functional TCR sequences in their endogenous format with desired characteristics. For the HERC1 neoantigen, a hybrid simulation- and AI-based approach yielded two TCR candidates of high activity and mutant specificity, activating against reporter cells and killing cancer cells at levels beyond the best patient-derived TCR we could identify using long and extensive screening from blood. An AI-only approach showed more modest hit rates but was nonetheless successful in generating engineered TCRs against the EGFR T790M neoantigen.

TCR EC_750-5 showed encouraging activation and killing profiles, demonstrating more specific recognition of the cancer neoantigen relative to its wild-type peptide counterpart. Interestingly, the other EGFR-targeting TCR (EC_7-7) contained a cysteine residue in its CDR3β loop that may enable stronger activation and killing from strengthened peptide-TCR interactions. TCRs such as this are well characterized and can result in impressive TCR activation and strong ZAP70 signaling; TCRs with cysteines are considered for use in modulating TCR-pMHC interactions (*41*). It is important to note that this phenomenon has been observed in natural TCRs and is sometimes used in TCR engineering, but can be selected against during thymic selection due to risks of auto-immunity (*43*). In an engineering context, these structural motifs could either be avoided to prevent excessive signaling or strategically leveraged to enhance activation of weak binders, specifically in cases in which the neoantigen introduces a presented cysteine residue. These findings underscore the importance of incorporating structural and functional “rules” into design strategies to balance potency and safety. Future work may be need and should take these considerations in mind to optimize TCRs for clinical use.

While the modified RFdiffusion model used in this work as a point of comparison yielded intermediate TCR binding scores relative to our naïve and hybrid models, application of the most current all-atom diffusion models would undoubtedly be of interest in the context of endogenous TCR design. However, differences between de novo design use cases for antibodies and TCRs should be noted. One major advantage of diffusion models in antibody design is the ability to target arbitrary epitopes or regions of interactions that might be informed by AI. In the context of TCRs, the space of possible epitopes is significantly restricted to the presented peptide, and concerns over specificity in this narrow interaction region might tend to take precedent over the novelty of binding modes. Additionally, minor changes to CDR-peptide interactions at the immune synapse can have major implications in determining both binding amplitude and specificity, a constraint that antibody generative models have not necessarily had to prioritize outside of MHC-peptide contexts.

Regardless of the specific AI-guided workflow, physics-based simulations proved useful in both validating and prioritizing TCR candidates, showing particularly accurate results with the HERC1 neoantigen as design components. The integration of physics-based considerations represents a major topic of interest for continued AI model development and should be considered to facilitate the intricate needs of TCR design against cancer neoantigens.

It is tantalizing to speculate that our *in-silico* TCR engineering strategy resembles a form of AI-driven biomimicry, adapting the principles of antibody maturation, where antibodies evolve through iterative mutation and selection, to TCR design. Unlike antibodies, which naturally undergo somatic hypermutation to improve binding, TCRs lack such a refinement process in organisms. By combining free-energy perturbation (FEP) simulations with generative AI, we effectively recreate maturation cycles computationally: predicting beneficial mutations, recombining them into multi-mutant candidates, and iteratively prioritizing sequences for improved binding and specificity. This approach addresses the unique constraints of TCR design for neoantigens, where the TCR must discriminate between mutant and wild-type peptides differing by a single residue while engaging two distinct surfaces, the peptide and the MHC. Minor changes in CDR–peptide interactions can dramatically alter binding amplitude and specificity, making this a uniquely challenging design problem. This work sets the stage for a new strategy in immunotherapy, one where TCRs can be rationally designed with the same precision and scalability that has propelled antibodies to the forefront of biologics.

## Acknowledgements

We first thank each patient and their families for participating in this study. We thank members of the Chan lab for expert advice and collaboration. We also thank Jianying Hu, Michal Rosen-Zvi, and Ajay Royyuru for insightful comments. We thank Drew Pardoll for providing advice on TCR assays.

## Supplementary Materials

### Data and Materials Availability

Sequencing, flow cytometry, simulation, and AI modeling files will be available upon request.

### Materials and Methods

#### Cell culture

NCI-H1975 (ATCC CRL-5908) and TCR Knockout (KO) Jurkat cells (BPS Bioscience #78539) and all derivative lines were maintained in RPMI-1640 medium (Cleveland Clinic Media Core) supplemented with 10% v/v fetal bovine serum (FBS, Gibco #A52567-01) and 1% v/v penicillin-streptomycin (Pen-Strep, Gibco #15140122). K562 A*11:01 and T2 A*02:05 were maintained in IMDM with GlutaMAX (Gibco #31980030) supplemented with 10% v/v FBS and 1% v/v Pen-Strep. Lenti-X 293T (Takara #632180) and HEK293-GP (Takara #631458) cells were maintained in DMEM (Cleveland Clinic Media Core) supplemented with 10% v/v FBS and 1% v/v Pen-Strep. All cell lines were cultured at 37 °C in a humidified incubator with 5% CO_2_ and were confirmed to be mycoplasma-negative throughout the study.

PBMCs were isolated from de-identified, healthy donor leukopaks (New York Blood Center) by Ficoll density centrifugation (Cytiva #17544602). Briefly, leukopaks were diluted 1:1 ratio with RPMI media and carefully dispensed onto 10mL of Ficoll-Paque density gradient media, split across two 50mL tubes. Tubes were spun at 400 × g for 20min at 25°C with brake off. Buffy coat layer containing PBMCs was collected, washed 2x with 50mL RPMI and spun 400 × g for 10min at 25°C before counting and cryopreservation with CTL-Cryo ABC (CTLC-ABC), following manufacturer’s instructions.

#### Generation of reporter and stable cell lines

TCR KO Jurkat NFAT reporter cells were generated by transduction with retrovirus encoding 8xNFAT-ZsG-hCD8 (Addgene #153417) followed by FACS sorting for hCD8 surface expression. K562 cells were nucleofected with HLA-A*11:01 cloned into pcDNA3.1(-) and selected with geneticin (G418) for 14 days. Top-positive clones were isolated by single cell sorting and maintained in geneticin (1µg/mL). T2-A*02:05 cells were generated by CRISPR-Cas9 knockout of endogenous HLA-A*02:01 in parental T2 cells followed by electroporation with a mammalian expression vector containing HLA-A*02:05 using the Lonza 4D Nucleofector. Cells with highest regained HLA-A2 (clone BB.2) expression were isolated by FACS sorting and expanded in culture. NCI-H1975 A*02:05 was made by lentiviral transduction of pLV-HLA-A*02:05-Hygro into H1975 followed by hygromycin selection. All stable cell lines were confirmed by flow cytometry for relevant surface expression before being used in experiments.

#### Design of TCR constructs

TCR alpha and beta variable regions were previously retrieved from single-cell V(D)J sequencing (iRepertoire and 10x Genomics) of antigen-specific stimulated T cells (*1*). Reconstruction of full-length TCRs was performed as previously described(*2*). Briefly, TRAV and TRBV were fused to mouse TRA and TRB constant regions. The murine constant regions were modified to introduce an additional disulfide bond and with additional transmembrane hydrophobic residues to allow preferential pairing (*3, 4*). TCR beta and alpha chains were separated by a Furin-P2A linker, codon optimized and synthesized as gBlocks (Twist Biosciences) and subsequently cloned into the pMSGV retroviral backbone (Addgene #128176) between PacI and NotI restriction site with HiFi assembly (NEB #E2621).

For primary TCR-T cell production, TCRs of interest were subcloned with HiFi assembly into a lentiviral backbone pTRPE (gift of Jos Melenhorst lab), which contains a HIV Ψ packaging element, Rev-responsive element, cPPT/CTS, EF-1α promoter, and woodchuck hepatitis virus post-transcriptional regulatory element, flanked by chimeric 5’ long terminal repeats (LTR) and 3’ ΔU3 LTR.

#### Retrovirus and Lentivirus production

Retroviral particles encoding TCR constructs were produced by co-transfection of HEK293-GP cells with pMSGV-TCR and pCMV-VSV-G (Addgene #8454) plasmids at 2:1 ratio using Lipofectamine 3000 (ThermoFisher #L3000075). Retroviral supernatant was harvested after 48 and 72 hours post-transfection and concentrated using Retro-X concentrator (Takara #631455). Similarly, lentiviral particles containing the TCRs were produced by co-transfection of Lenti-X 293T cells with 6 µg pTRPE-Rev, 2.33 µg pTRPE-VSVg, 6 µg pTRPE-Gag-Pol, and 5 µg pTRPE-TCR with 29 µL of Lipofectamine 3000 in a 10cm dish. Lentiviral supernatant was harvested at 48 and 72 hours post-transfection and concentrated using Lenti-X concentrator (Takara #631232). Concentrated viral particles were resuspended in PBS and stored at -80°C until use.

#### HLA stabilization assay

Peptides were synthesized by Genscript at a purity >90%, reconstituted in DMSO (Sigma #D2650) to 50mM, aliquoted, and stored at –80°C. T2 A*02:05 were loaded with peptides (10µM) and incubated in culture medium at 37°C. After 16 hrs, cells were analyzed for HLA-ABC and β2-microglobulin (B2M) expression by flow cytometry.

#### Primary T cell culture and lentiviral transduction

Healthy donor PBMCs were resuspended in PBS supplemented with 2% FBS and 1 mM EDTA. CD3+ T cells were isolated with untouched negative selection using EasySep Human T cell Isolation Kit (STEMCELL Technologies #17951). Purified CD3+ T cells were cultured in IMDM supplemented with 10% FBS, 50U/mL IL-2 (Peprotech #200-02), 5ng/mL IL-15 (Peprotech #200-15), 5ng/mL IL-7 (Peprotech #200-07) and stimulated with Dynabeads® Human T-Expander CD3/CD28 (ThermoFisher #11132D) at a bead-to-cell ratio of 3:1 for 48 hours prior to transduction. For transduction of TCRs by spinoculation, 100 µL of concentrated lentivirus was added to a Retronectin (Takara #T100B) coated 24-well plate (10 µg/cm^2^) and centrifuged at 2000 × g for 2 hours at 32 °C. Virus-coated wells were washed with 2% BSA in PBS. Next, T cells were resuspended at 5 × 10^5^ cells/mL in IMDM + 10% FBS supplemented with 200 U/mL IL-2, and 1 mL of cells was added per well. Plates were centrifuged at 1500 rpm (∼532 × g) for 15 minutes at 32 °C and then transferred to a 37 °C incubator. Transduction efficiency was assessed by flow cytometry 72-96 hours post-infection, and cells were used after 96 hours post-transduction.

#### T cell activation assays

K562 cells expressing HLA-A*11:01 were pulsed with HERC1 peptides (10µM) for 2 hrs at 37°C. Cells were pelleted, washed, and resuspended in fresh culture medium without peptide. Jurkats were then added at an effector:target (E:T) ratio of 2:1 and co-cultured at 37 °C. After 16 h, cells were harvested and analyzed for surface activation markers and ZsGreen expression by flow cytometry. For intracellular cytokine analysis, primary T cells and K562 A*11:01 were co-cultured in the presence of brefeldin A (2× final concentration, per manufacturer’s recommendation, Biolegend #420601). After 16 h, cells were permeabilized with the Fixation/Permeabilization kit (ThermoFisher #00-5521-00) and stained for intracellular cytokines using anti-TNFα and anti-IFNγ antibodies and analyzed by flow cytometry.

#### xCELLigence killing assays

Tumor target cell lines (K562 A*11:01 or NCI-H1975 A*02:05) were plated onto E-Plate 96 PET (Agilent #300600910) plates at 10,000 cells/well and allowed to adhere overnight. After 16-20 hours, 1 × 10^6^ primary T cells (10:1 E:T ratio) expressing the engineered or patient/donor-derived TCRs were added to the plate. Tumor only and full lysis (with 1% v/v Triton-X-100) controls were included in the experimental design. Cytotoxicity of the TCR-T cells was evaluated using xCELLigence RTCA MP (Agilent), which measures change in impedance due to adherent cells in each well every 15 minutes until end of the experiment. The data were processed using the xCELLigence RTCA software, and results reported as Cell Index, normalized to time when T cells were added, or Percent Cytolysis, which is calculated by the following formula:

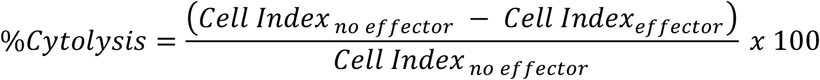

For K562 killing assays, E-Plates were coated with anti-CD71 antibodies one day prior to allow for K562 tumor cell adherence to plate using the anti-CD71 tethering kit (Agilent #8100017) following manufacturer’s instructions. 16 hours later, K562s were loaded with 10µM peptide for 2 hours before 100,000 primary T cells expressing engineered TCRs or non-transduced control (NTC) T cells were added to each well (10:1 E:T ratio). Tumor only and full lysis with 2% Triton X-100 controls were included in the experimental design.

#### Molecular Dynamics and Free Energy Perturbation Simulations

Physics-based simulations (referring to both molecular dynamics (MD) simulations and free-energy perturbation calculations (FEP)) were conducted via a typical procedure using NAMD3(*5*) and the CHARMM36m force field(*6*). For molecular dynamics on the HERC1 mutant and wildtype complexes, starting structures were created based on a template PDB structure (PDB: 5NMF)(*7*) using standard homology modeling software(*8*) and the MUTATOR plugin in VMD (*9*), minimized with 10,000 steepest descent steps, equilibrated in the NPT ensemble with harmonic constraints, and then run at production for 200-500 ns without constraints at 310K in the NPT ensemble. Initial components of complex structures (MHC-peptide complexes and TCR structures) were generated with AlphaFold2-multimer (*10*) and minimized as described above. In the case of the HERC1 patient-derived reference TCR, initial simulations were annealed with two minimization runs following 5 ns of constrained equilibration. For EGFR TCRs, the optimized HERC1 reference structure was then used as a starting structure for alignment and to seed minimization, and the above procedure was repeated as described.

Input structures for alchemical FEP simulations were created with a similar approach to that described above, with hybrid residues being adapted from the CHARMM36m force field. Systems were also minimized and equilibrated as described above, and production simulations were run through alchemical lambda space with windows spaced at 0.04 outside of finer creation and annihilation regions. Softcore potentials and electrostatic decoupling were carried out in a standard fashion. Individual simulation runs were replicated until random errors (95% confidence intervals) fell below 2.0 kcal/mol for ΔΔG values.

Candidates were enumerated and prioritized based on favorability of FEP scores in single and multiple mutant iterations, respectively. Single mutation FEP results were used to identify favorable points in the CDR3β sequence for candidate enumeration: point mutations that yielded negative ΔΔG predictions were curated and inserted into sequences in all possible double, triple, and quadruple mutant configurations, yielding a total of 9061 candidates for advancement to graph-based AI surrogates. After extracting top multiple mutant candidates from the graph-based AI surrogate pipeline described below, those TCRs were subjected to FEP for final ranking.

#### Graph-Based AI Surrogates for Free Energy Change Prediction

Structure-based AI models trained to predict binary labels for favorable (negative ΔΔG) interactions between mutant TCR-pMHC complexes constructed using a 3D graph-based architecture from previous work (*11*). Internal representations for mutant and reference complexes were subtracted to yield a representation vector on which model loss functions were minimized. Models were pre-trained on all available single-point FEP data for the HERC1 system, with input complex structures being generated with VMD Mutator and aligned to reference TCR-pMHC complexes prior to graph featurization and input into the graph convolutional model encoder.

Graph embedding vectors were collapsed with pooling operations and subjected to a softmax layer to yield multiple mutant candidates scoring in the 95^th^ percentile or above were clustered using a standard k-means approach, and eight cluster centers were advanced to final validation with free energy simulations and LLM-based predictive models.

#### LLM-Based AI Models for Binding Prediction and Candidate Generation

To facilitate LLM-based components of our TCR design pipeline, the MAMMAL (Molecular Aligned Multi-Modal Architecture and Language) foundation model was applied, as described in previous work (*12*). MAMMAL is a sequence-to-sequence cross-modal foundation model pre-trained on over 2 billion samples across protein and antibody sequences, small molecules, and gene expression profiles. It supports a range of tasks, including classification, regression, and generation, through a hybrid encoder-decoder architecture that integrates sequence-based representations of diverse biomedical modalities. This framework enables effective modeling of complex interactions, such as those at the pMHC–TCR interface, by leveraging structured prompts that combine tokens and scalar values for precise predictions and de-novo sequence generation.

For TCR-specific applications in this study, MAMMAL was further pre-trained on an unmasking task on a large-scale dataset of TCR CDR3β sequences derived from the NCBI Sequence Read Archive (SRA) (*13*), comprising 126,432,303 unique CDR3 sequences. Data extraction from NCBI SRA was performed using the SRA Toolkit https://github.com/ncbi/sra-tools, which enabled efficient downloading and processing of RNA-seq datasets. Nucleotide sequences were subsequently translated to CDR3 amino acid sequences using Decombinator v2.2 software (*14*). Additionally, the original MAMMAL model was fine-tuned on the Weber TCR binding dataset (also known as TITAN) (*15*) for TCR Beta chain-epitope binding prediction; for full details, see Shoshan et al.(*12*). This additional pre-training enhanced the model’s ability to generate and rank CDR3β variants tailored to neoantigen binding, improving its alignment with the natural diversity and functional constraints of TCR repertoires.

In the hybrid physics-AI approach, a version of MAMMAL further fine-tuned for pMHC-TCR binding prediction was employed. Specifically, the model was adapted to predict TCR– neoantigen binding affinities, guiding the selection and refinement of candidates generated from physics-based simulations.

In the pure-AI workflow, two complementary approaches were applied for candidate generation. First, a “naive” method involved infilling CDR3β region conditioned on the target epitope, directly leveraging the model’s generative capabilities to produce sequences optimized for binding. Second, we used an iterative multiple constraint generation method, Ratner V, Shoshan Y, Raboh M, Gurev V, Golts A, Weber J, Hexter E., US Patent Pending, to produce candidates that exhibit stronger predicted binding to mutated epitopes compared to wild-type epitopes. This approach combines inference scores from multiple targets (e.g., mutant vs. wild-type epitopes) to guide sequence unmasking, ensuring specificity while optimizing for desired interactions.

#### Diffusion-Based AI Models for Candidate Generation

To produce diffusion AI-based candidates as a point of comparison, RFdiffusion was applied using default backbone generation parameters. A modified version of ProteinMPNN was used for the explicit sequence generation/inverse folding step: custom positional contraints were applied based on relative residue frequencies in CDR regions within the SAbDab(*16*) database and further conditioned on patient-derived reference sequences with an adjustment of 0.2 units at each native position. A total of 16 sequences were created for each of the 16 backbone variations generated, and duplicate sequences were removed prior to analysis.

## Supplemental Figures

**Supplemental Figure 1:**
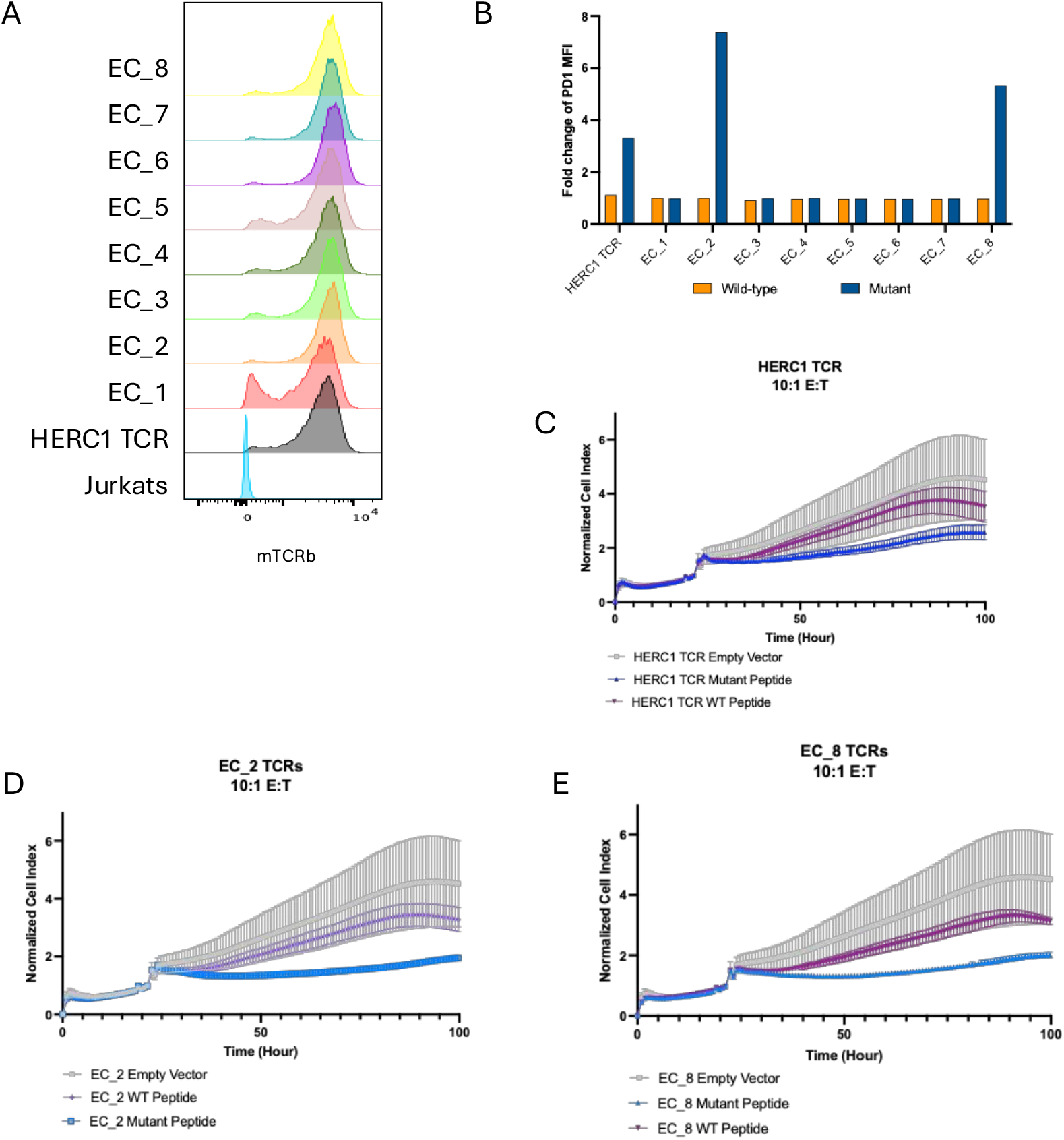
(**A**) Histograms showing mTCRb surface expression levels in Jurkats after transduction of retrovirus carrying HERC1 engineered TCRs (**B**) hCD8-NFAT-Jurkats were transduced with HERC1 TCRs and AI-designed TCRs. Activation of TCRS by HLA-A*11:01-K562s pulsed with 10 μM of WT HERC1 and MT HERC1 peptides as shown by PD1 expression. MFI of protein was normalized to DMSO control to calculate fold change. (**C-E**) HLA-A*11:01-K562s were tethered to E-plate for 2hrs at RT, and TCRS were added at E:T ratio of 10:1. T cell–mediated Killing was monitored using the xCELLigence Real-Time Cell Analyzer over 100hr. Cell index is plotted as a measure of total live tumor cells over time. The result shown is representative of 2 independent experiments.

**Supplemental Fig. 2.**
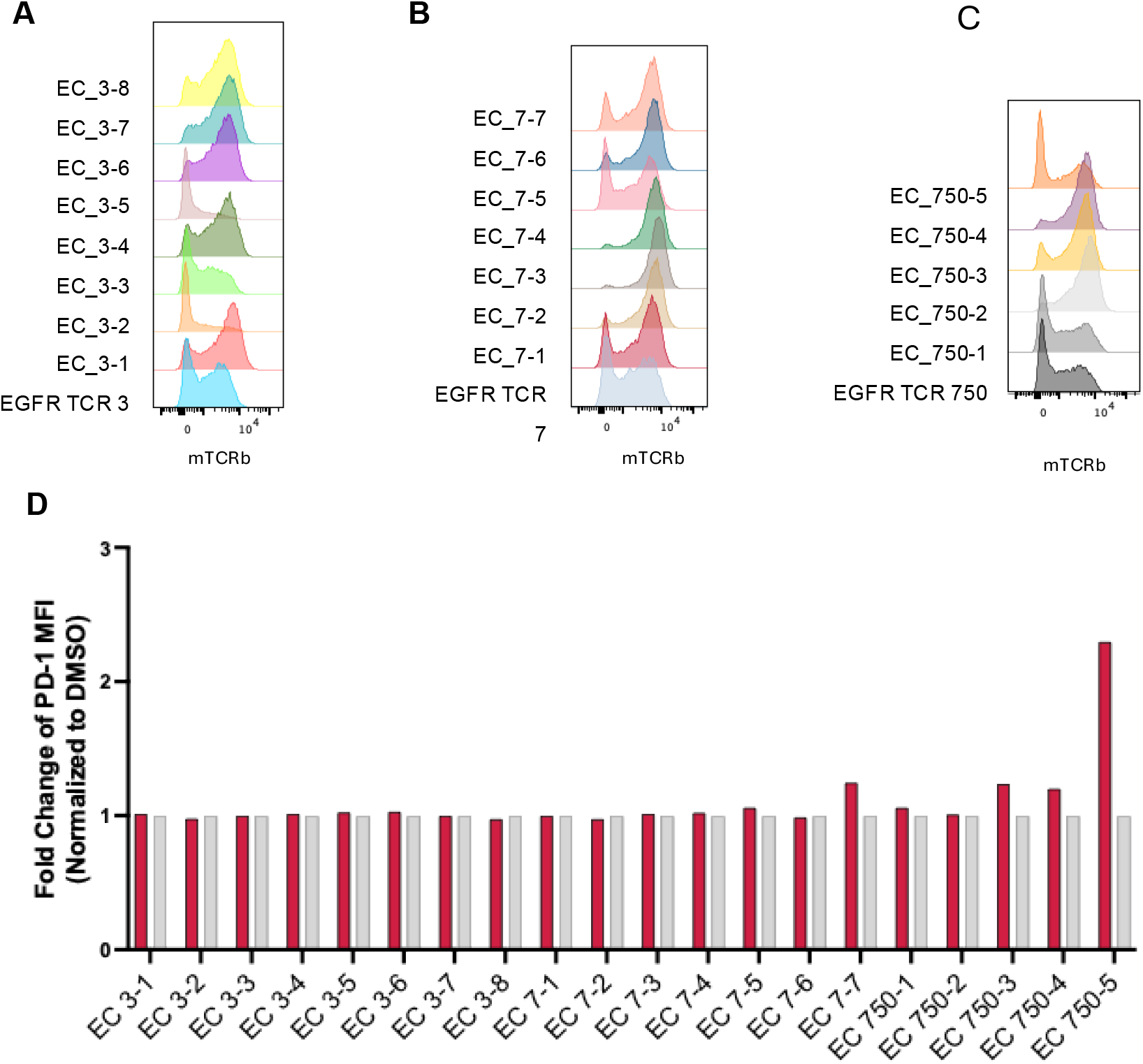
**(A-C)** Histograms showing mTCRb surface expression levels in Jurkats after transduction of retrovirus carrying EGFR engineered TCRs (**D**)PD-1 surface expression on Jurkat reporter cells during activation co-culture assays with HLA-A*02:05-expressing APCs pulsed with 10μM EGFR neopeptide. MFI of PD-1 expression was normalized to DMSO vehicle control to calculate fold change.

**Supplemental Fig. 3.**
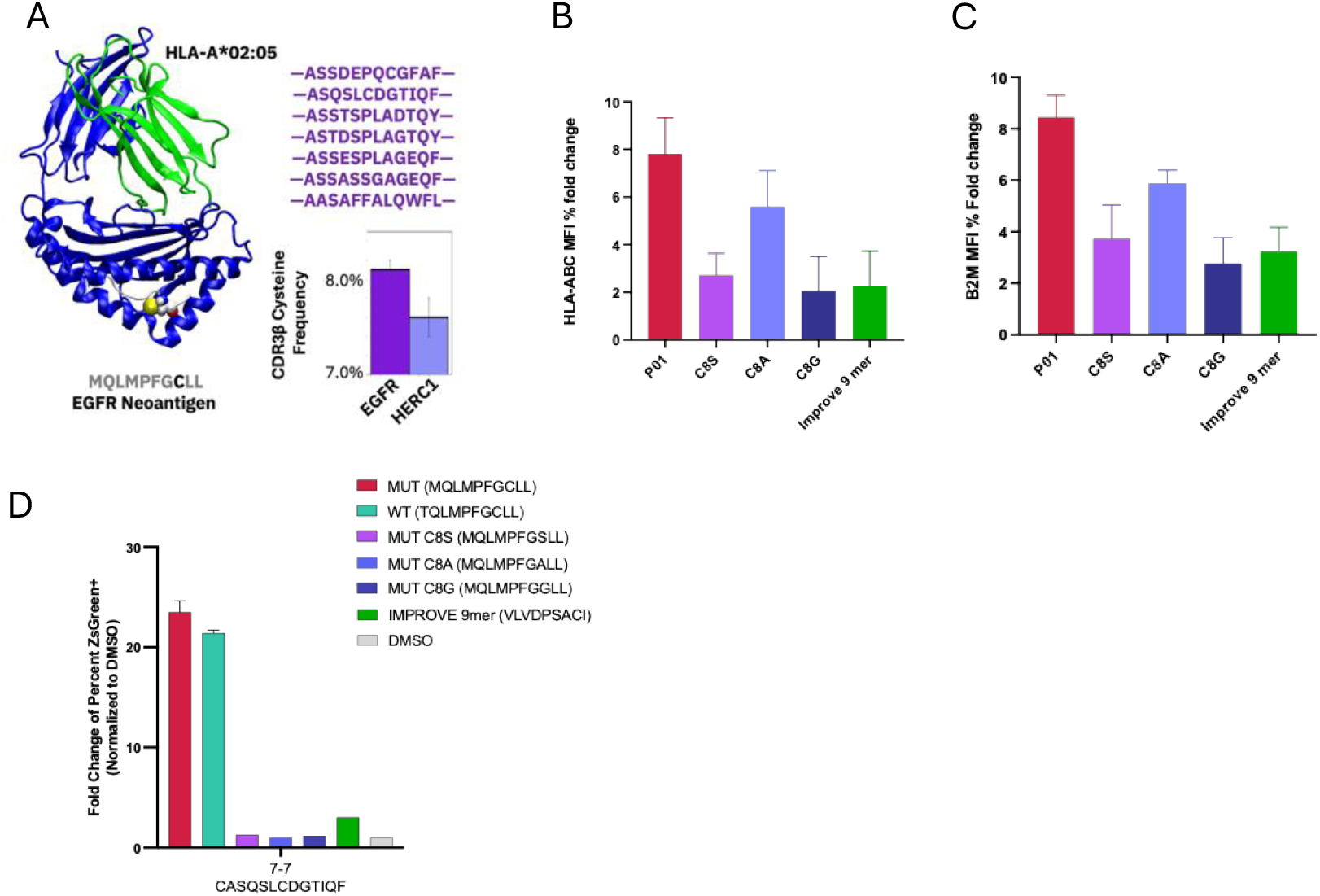
**(A-D)**. (A) TCR candidate CDR3β sequences generated with the pure-AI approach against the EGFR neoantigen (MQLMPFGCLL) and cysteine frequency comparison plot between HERC1 and EGFR candidates. (B-C) HLA stabilization assay showing surface expression MFI of HLA and beta-2-microglobulin (β2M) after peptide stimulation. (D) Activation assay of EC_7-7 with 10μM of mutated control peptides C8A, C8G, C8S and improve database neoantigen with P8 cysteine. Error bars display standard deviation calculated from technical triplicates.

## References

1. A. Desrichard, A. Snyder, T. A. Chan, Cancer Neoantigens and Applications for Immunotherapy. Clin Cancer Res 22, 807–812 (2016).

2. P. Team et al., PXDesign: Fast, Modular, and Accurate De Novo Design of Protein Binders. bioRxiv, 2025.2008.2015.670450 (2025).

3. T. J. Alban et al., Neoantigen immunogenicity landscapes and evolution of tumor ecosystems during immunotherapy with nivolumab. Nat Med 30, 3209–3222 (2024).

4. E. H. Hsiue et al., Targeting a neoantigen derived from a common TP53 mutation. Science 371, (2021).

5. T. T. Spear, B. D. Evavold, B. M. Baker, M. I. Nishimura, Understanding TCR affinity, antigen specificity, and cross-reactivity to improve TCR gene-modified T cells for cancer immunotherapy. Cancer Immunol Immunother 68, 1881–1889 (2019).

6. S. Romagnani, Immunological tolerance and autoimmunity. Intern Emerg Med 1, 187–196 (2006).

7. M. G. Weigert, I. M. Cesari, S. J. Yonkovich, M. Cohn, Variability in the lambda light chain sequences of mouse antibody. Nature 228, 1045–1047 (1970).

8. K. D. Householder et al., De novo design and structure of a peptide-centric TCR mimic binding module. Science 389, 375–379 (2025).

9. A. Madani et al., Large language models generate functional protein sequences across diverse families. Nat Biotechnol 41, 1099–1106 (2023).

10. B. L. Hie et al., Efficient evolution of human antibodies from general protein language models. Nat Biotechnol 42, 275–283 (2024).

11. A. Elnaggar et al., ProtTrans: Toward Understanding the Language of Life Through Self-Supervised Learning. IEEE Trans Pattern Anal Mach Intell 44, 7112–7127 (2022).

12. K. H. Johansen et al., De novo-designed pMHC binders facilitate T cell-mediated cytotoxicity toward cancer cells. Science 389, 380–385 (2025).

13. A. Motmaen et al., Targeting peptide-MHC complexes with designed T cell receptors and antibodies. bioRxiv, (2025).

14. D. Nori, S. V. Mathis, A. Shanehsazzadeh, Evaluating Zero-Shot Scoring for In Vitro Antibody Binding Prediction with Experimental Validation. arXiv preprint 2312.05273, (2023).

15. N. Bio, S. Biswas, *De novo* design of epitope-specific antibodies against soluble and multipass membrane proteins with high specificity, developability, and function. bioRxiv, 2025.2001.2021.633066 (2025).

16. Y. Bang et al., Precise, Specific, and Sensitive *De Novo* Antibody Design Across Multiple Cases. bioRxiv, 2025.2003.2009.642274 (2025).

17. N. R. Bennett et al., Atomically accurate de novo design of antibodies with RFdiffusion. bioRxiv, (2025).

18. C. D. Team et al., Zero-shot antibody design in a 24-well plate. bioRxiv, 2025.2007.2005.663018 (2025).

19. A. Shanehsazzadeh et al., IgDesign: <em>In vitro</em> validated antibody design against multiple therapeutic antigens using inverse folding. bioRxiv, 2023.2012.2008.570889 (2024).

20. J. L. Watson et al., De novo design of protein structure and function with RFdiffusion. Nature 620, 1089–1100 (2023).

21. N. A. Rizvi et al., Cancer immunology. Mutational landscape determines sensitivity to PD-1 blockade in non-small cell lung cancer. Science 348, 124–128 (2015).

22. T. Yamada et al., EGFR T790M mutation as a possible target for immunotherapy; identification of HLA-A*0201-restricted T cell epitopes derived from the EGFR T790M mutation. PLoS One 8, e78389 (2013).

23. J. C. Phillips et al., Scalable molecular dynamics on CPU and GPU architectures with NAMD. J Chem Phys 153, 044130 (2020).

24. J. Huang et al., CHARMM36m: an improved force field for folded and intrinsically disordered proteins. Nat Methods 14, 71–73 (2017).

25. D. K. Cole et al., Dual Molecular Mechanisms Govern Escape at Immunodominant HLA A2-Restricted HIV Epitope. Front Immunol 8, 1503 (2017).

26. A. Fiser, A. Sali, Modeller: generation and refinement of homology-based protein structure models. Methods Enzymol 374, 461–491 (2003).

27. W. Humphrey, A. Dalke, K. Schulten, VMD: visual molecular dynamics. J Mol Graph 14, 33-38, 27-38 (1996).

28. P. Bryant, G. Pozzati, A. Elofsson, Improved prediction of protein-protein interactions using AlphaFold2. Nat Commun 13, 1265 (2022).

29. J. K. Weber et al., Unsupervised and supervised AI on molecular dynamics simulations reveals complex characteristics of HLA-A2-peptide immunogenicity. Brief Bioinform 25, (2023).

30. D. Chowell et al., Patient HLA class I genotype influences cancer response to checkpoint blockade immunotherapy. Science 359, 582–587 (2018).

31. Y. Shoshan et al., MAMMAL--Molecular Aligned Multi-Modal Architecture and Language. arXiv preprint 2410.22367, (2024).

32. Y. Kodama, M. Shumway, R. Leinonen, C. International Nucleotide Sequence Database, The Sequence Read Archive: explosive growth of sequencing data. Nucleic Acids Res 40, D54–56 (2012).

33. N. Thomas, J. Heather, W. Ndifon, J. Shawe-Taylor, B. Chain, Decombinator: a tool for fast, efficient gene assignment in T-cell receptor sequences using a finite state machine. Bioinformatics 29, 542–550 (2013).

34. A. Weber, J. Born, M. Rodriguez Martinez, TITAN: T-cell receptor specificity prediction with bimodal attention networks. Bioinformatics 37, i237–i244 (2021).

35. J. Dunbar et al., SAbDab: the structural antibody database. Nucleic Acids Res 42, D1140–1146 (2014).

36. J. Dauparas et al., Robust deep learning-based protein sequence design using ProteinMPNN. Science 378, 49–56 (2022).

37. D. del Alamo, R. Frick, D. Truan, J. Karpiak, Adapting ProteinMPNN for antibody design without retraining. bioRxiv, 2025.2005.2009.653228 (2025).

38. L. McInnes, J. Healy, J. Melville, Umap: Uniform manifold approximation and projection for dimension reduction. arXiv preprint 1802.03426, (2018).

39. R. C. Wirasinha et al., αβ T-cell receptors with a central CDR3 cysteine are enriched in CD8αα intraepithelial lymphocytes and their thymic precursors. Immunology and Cell Biology 96, 553–561 (2018).

40. A. M. Rosenberg, C. M. Ayres, A. V. Medina-Cucurella, T. A. Whitehead, B. M. Baker, Enhanced T cell receptor specificity through framework engineering. Front Immunol 15, 1345368 (2024).

41. C. Szeto et al., Covalent TCR-peptide-MHC interactions induce T cell activation and redirect T cell fate in the thymus. Nat Commun 13, 4951 (2022).

42. A. Borch et al., IMPROVE: a feature model to predict neoepitope immunogenicity through broad-scale validation of T-cell recognition. Front Immunol 15, 1360281 (2024).

43. S. R. Daley et al., Cysteine and hydrophobic residues in CDR3 serve as distinct T-cell self-reactivity indices. J Allergy Clin Immunol 144, 333–336 (2019).

## Supplemental References

1. N. A. Rizvi et al., Cancer immunology. Mutational landscape determines sensitivity to PD-1 blockade in non-small cell lung cancer. Science 348, 124–128 (2015).

2. S. Stevanovic et al., Landscape of immunogenic tumor antigens in successful immunotherapy of virally induced epithelial cancer. Science 356, 200–205 (2017).

3. C. J. Cohen et al., Enhanced antitumor activity of T cells engineered to express T-cell receptors with a second disulfide bond. Cancer Res 67, 3898–3903 (2007).

4. A. Haga-Friedman, M. Horovitz-Fried, C. J. Cohen, Incorporation of transmembrane hydrophobic mutations in the TCR enhance its surface expression and T cell functional avidity. J Immunol 188, 5538–5546 (2012).

5. J. C. Phillips et al., Scalable molecular dynamics on CPU and GPU architectures with NAMD. J Chem Phys 153, 044130 (2020).

6. J. Huang et al., CHARMM36m: an improved force field for folded and intrinsically disordered proteins. Nat Methods 14, 71–73 (2017).

7. D. K. Cole et al., Dual Molecular Mechanisms Govern Escape at Immunodominant HLA A2-Restricted HIV Epitope. Front Immunol 8, 1503 (2017).

8. A. Fiser, A. Sali, Modeller: generation and refinement of homology-based protein structure models. Methods Enzymol 374, 461–491 (2003).

9. W. Humphrey, A. Dalke, K. Schulten, VMD: visual molecular dynamics. J Mol Graph 14, 33-38, 27-38 (1996).

10. P. Bryant, G. Pozzati, A. Elofsson, Improved prediction of protein-protein interactions using AlphaFold2. Nat Commun 13, 1265 (2022).

11. J. K. Weber et al., Unsupervised and supervised AI on molecular dynamics simulations reveals complex characteristics of HLA-A2-peptide immunogenicity. Brief Bioinform 25, (2023).

12. Y. Shoshan et al., MAMMAL--Molecular Aligned Multi-Modal Architecture and Language. arXiv preprint 2410.22367, (2024).

13. Y. Kodama, M. Shumway, R. Leinonen, C. International Nucleotide Sequence Database, The Sequence Read Archive: explosive growth of sequencing data. Nucleic Acids Res 40, D54–56 (2012).

14. N. Thomas, J. Heather, W. Ndifon, J. Shawe-Taylor, B. Chain, Decombinator: a tool for fast, efficient gene assignment in T-cell receptor sequences using a finite state machine. Bioinformatics 29, 542–550 (2013).

15. A. Weber, J. Born, M. Rodriguez Martinez, TITAN: T-cell receptor specificity prediction with bimodal attention networks. Bioinformatics 37, i237–i244 (2021).

16. J. Dunbar et al., SAbDab: the structural antibody database. Nucleic Acids Res 42, D1140–1146 (2014).

